# The Preprocessed Connectomes Project Repository of Manually Corrected Skull-stripped T1-weighted Anatomical MRI Data

**DOI:** 10.1101/067017

**Authors:** Benjamin Puccio, James P Pooley, John S Pellman, Elise C Taverna, R Cameron Craddock

## Abstract

**Background:** Skull-stripping is the procedure of removing non-brain tissue from anatomical MRI data. This procedure is necessary for calculating brain volume and for improving the quality of other image processing steps. Developing new skull-stripping algorithms and evaluating their performance requires gold standard data from a variety of different scanners and acquisition methods. We complement existing repositories with manually-corrected brain masks for 125 T1-weighted anatomical scans from the Nathan Kline Institute Enhanced Rockland Sample Neurofeedback Study.

**Findings:** Skull-stripped images were obtained using a semi-automated procedure that involved skull-stripping the data using the brain extraction based on non local segmentation technique (BEaST) software and manually correcting the worst results. Corrected brain masks were added into the BEaST library and the procedure was reiterated until acceptable brain masks were available for all images. In total, 85 of the skull-stripped images were hand-edited and 40 were deemed to not need editing. The results are brain masks for the 125 images along with a BEaST library for automatically skull-stripping other data.

**Conclusion:** Skull-stripped anatomical images from the Neurofeedback sample are available for download from the Preprocessed Connectomes Project. The resulting brain masks can be used by researchers to improve their preprocessing of the Neurofeedback data, and as training and testing data for developing new skull-stripping algorithms and evaluating the impact on other aspects of MRI preprocessing. We have illustrated the utility of these data as a reference for comparing various automatic methods and evaluated the performance of the newly created library on independent data.

## Data Description

One of the many challenges facing the analysis of magnetic resonance imaging (MRI) data is achieving accurate brain extraction from the data. Brain extraction, also known as skull-stripping, aims to remove all nonbrain tissue from an image. This is one of the preliminary steps in preprocessing and the quality of its result affects the subsequent steps, such as image registration and brain matter segmentation. There are a multitude of challenges that surround the process of brain extraction. The manual creation and correction of brain masks is tedious, time-consuming, and susceptible to experimenter bias. On the other hand, fully automated brain extraction is not a simple image segmentation problem. Brains in images can differ in orientation and morphology, especially in pediatric, geriatric, and pathological brains. In addition, non-brain tissue may resemble brain in terms of voxel intensity. Differences in MRI scanner, acquisition sequence, and scan parameters can also have an effect on automated algorithms due to differences in image contrast, quality, and orientation. Image segmentation techniques with low computational time, high accuracy, and high flexibility are extremely desirable.

Developing new automated skull-stripping methods, and comparing these with existing methods, requires large quantities of *gold standard* skull-stripped data acquired from a variety of scanners using a variety of sequences and parameters. This is due to the variation in performance of algorithms using different MRI data. Repositories containing gold standard skull-stripped data already exist: The Alzheimer’s Disease Neuroimaging Initiative (ADNI) [1]; BrainWeb: Simulated Brain Database (SBD) [2]; The Internet Brain Segmentation Repository (IBSR) at the Center for Morphometric Analysis [3]; the LONI Probabilistic Brain Atlas (LPBA40) at the UCLA Laboratory of Neuro Imaging [4]; and The Open Access Series of Imaging Studies (OASIS) [5], the last of which is not manually delineated but has been used as gold standard data [6, 7]. We extend and complement these existing repositories by releasing manually corrected skull strips for 125 individuals from the NKI Enhanced Rockland Sample Neurofeedback study.

### Data acquisition

The repository was constructed from defaced and anonymized anatomical data downloaded from the Nathan Kline Institute Enhanced Rockland Sample Neurofeedback Study (NFB) [?]. The NFB is a 3-visit study that involves a deep phenotypic assessment on the first and second visits [?], a 1-hour connectomic MRI scan on the second visit, and a 1-hour neurofeedback scan on the last visit. Up to 3 months may have passed between the first and last visits. The 125 participants included 77 females and 48 males in the 21–45 age range (average: 31, standard deviation: 6.6). Sixty-six (66) of the participants had one or more current or past psychiatric diagnosis as determined by the structured clinical interview for the DSM IV (SCID) [8] [see Table 1]. No brain abnormalities or incidental findings were present in the included images as determined by a board-certified neuroradiologist. None of the participants had any other major medical condition such as cancer or AIDS. All experimental procedures were performed with institutional review board approval and only after informed consent was obtained.

**Table 1.** Neurofeedback Participant Diagnoses.

| Diagnosis (SCID #) | # |
| --- | --- |
| No Diagnosis or Condition on Axis I | 59 |
| Major Depressive Disorder, Past | 26 |
| Alcohol Abuse, Past (305) | 21 |
| Cannabis Abuse, Current | 11 |
| Cannabis Dependence, Past | 11 |
| Attention-Deficit/Hyperactivity Disorder, Current | 10 |
| Alcohol Dependence, Past | 5 |
| Posttraumatic Stress Disorder, Current | 5 |
| Specific Phobia, Past | 5 |
| Generalized Anxiety Disorder, Current | 4 |
| Cocaine Abuse, Past | 2 |
| Cocaine Dependence, Past | 2 |
| Hallucinogen Abuse, Past | 2 |
| Agoraphobia w/o History of Panic Disorder, Current | 2 |
| Anorexia Nervosa, Past | 2 |
| Anxiety Disorder NOS, Current | 2 |
| Panic Disorder w/ Agoraphobia, Past | 2 |
| Panic Disorder w/o Agoraphobia, Past | 2 |
| Social Phobia Current | 2 |
| Alcohol Abuse, Current | 1 |
| Amphetamine Dependence, Past | 1 |
| Bereavement | 1 |
| Body Dysmorphic Disorder, Current | 1 |
| Bulimia Nervosa, Current | 1 |
| Delusional Disorder Mixed Type | 1 |
| Eating Disorder NOS, Past | 1 |
| Hallucinogen Dependence, Past | 1 |
| Major Depressive Disorder, Current | 1 |
| Obsessive-Compulsive Disorder, Current | 1 |
| Opioid Abuse, Past | 1 |
| Phencyclidine Abuse, Past | 1 |
| Sedative, Hypnotic, or Anxiolytic Dependence, Past | 1 |
| Trichotillomania | 1 |

Anatomical MRI data from the third visit of the NFB protocol was used to build the Neurofeedback Skull-stripped (NFBS) repository. MRI data were collected on a 3 T Siemens Magnetom TIM Trio scanner (Siemens Medical Solutions USA: Malvern PA, USA) using a 12-channel head coil. Anatomical images were acquired at 1 × 1 × 1 mm^3^ resolution with a 3D T1-weighted magnetization-prepared rapid acquisition gradient-echo (MPRAGE) [9] sequence in 192 sagittal partitions each with a 256 × 256 mm^2^ field of view (FOV), 2600 ms repetition time (TR), 3.02 ms echo time (TE), 900 ms inversion time (TI), 8° flip angle (FA), and generalized auto-calibrating partially parallel acquisition (GRAPPA) acceleration [10] factor of 2 with 32 reference lines. The anatomical data was acquired immediately after a fast localizer scan and proceeded the collection of a variety of other scans [11], whose description is beyond the scope of this report.

### Brain Mask Definition

Many researchers differ on the standard for what to include and exclude from the brain. Some brain extraction methods, such as brainwash, include the dura mater in the brain mask to use as a reference for measurements [12]. The standard we used was adapted from Eskildsen et al (2012) [13]. Non-brain tissue is defined as skin, skull, eyes, dura mater, external blood vessels and nerves (e.g., optic chiasm, superior sagittal sinus, and transverse sinus). Cerebrum, cerebellum, brainstem, and internal vessels and arteries are included in the brain, along with cerebrospinal fluid (CSF) in ventricles, internal cisterns, and deep sulci.

### NFBS Repository Construction

The BEaST method (brain extraction based on nonlocal segmentation technique) was used to initially skull-strip the 125 anatomical T1-weighted images [13]. This software uses a patch-based label fusion method that labels each voxel in the brain boundary volume by comparing it to similar locations in a library of segmented priors. The segmentation technique also incorporates a multi-resolution framework in order to reduce computational time. The version of BEaST used was 1.15.00 and our implementation was based off of a shell script written by Qingyang Li [14]. The standard parameters were used in the configuration files and beast-library-1.1 was used for the initial skull-strip of the data. Before running mincbeast, the main segmentation script of BEaST, the anatomical images were normalized using the beast_normalize script. Mincbeast was run using the probability filter setting, which smoothed the manual edits, and the fill setting, which filled any holes in the masks. We found the failure rate for masks using BEaST was similar to that of the published rate of approximately 29% [13]. Visual inspection of these initial skull-stripped images indicated whether additional edits were necessary.

Manual edits were performed using the Freeview visualization tool from the FreeSurfer software package [?]. The anatomical image was loaded as a track volume and the brain mask was loaded as a volume. The voxel edit mode was then used to include or exclude voxels in the mask. As previously mentioned, all exterior non-brain tissue was removed from the head image, specifically the skull, scalp, fat, muscle, dura mater, and external blood vessels and nerves 1. Time spent editing each mask ranged from 1–8 hours, depending on the quality of the anatomical image and the BEaST mask. Afterwards, manually edited masks were used to populate the prior library of BEaST. This iterative bootstrapping technique was repeated until approximately 85 of the datasets were manually edited and all skull-strips were considered to be acceptable.

For each of the 125 subjects, the repository contains the de-faced and anonymized anatomical T1-weighted image, skull-stripped brain image, and brain mask. Each of these are in compressed NIfTI file format (.nii.gz). The size of the entire data set is around 1.9 GB.

**Figure 1.**
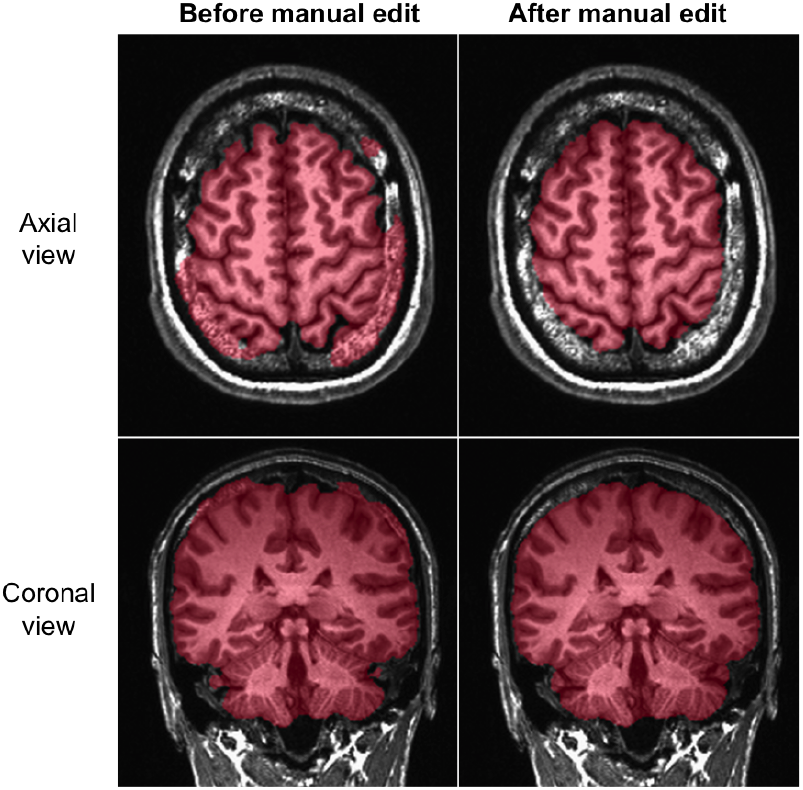
Manual Editing.

Axial and coronal slices in AFNI viewer of the brain mask and image pair, before and after manual editing in Freeview. The anatomical MRI image was loaded into the viewer as a grayscale image. The mask, which can be seen in a transparent red, was loaded as an overlay image.

### Data Validation

The semi-automated skull-stripping procedure was repeated until all brain masks were determined to be acceptable by two raters (BP and ET). Once this was completed, the brain masks were used as gold standard data for comparing different automated skull-stripping algorithms. Additionally, we evaluated the performance of the newly corrected BEaST library by comparing it to other skull-stripping methods on data from the Internet Brain Segmentation Repository (IBSR) [3] and LONI Probabilistic Brain Atlas (LPBA40) [4].

#### Skull-Stripping Algorithms

A wide variety of algorithms have been developed [6, 7, 12, 15, 16, 17, 18, 19], but we focused on FSL’s BET [20], AFNI’s 3dSkullStrip [21], and FreeSurfer’s Hybrid Watershed Algorithm (HWA) [22] based on their popularity.

- The *Brain Extraction Technique (BET)* is an algorithm incorporated in the FSL software that is based on a deformable model of the surface of the brain [20]. First, an intensity histogram is used to find the center of gravity of the head. Then a tessellated sphere is initialized around the center of gravity and expanded by locally adaptive forces. The method can also incorporate T2-weighted images to isolate the inner and outer skull and scalp. The bias field and neck setting (bet −B) was used since the anatomical images contained the subjects’ necks. The version of FSL that was used was 5.0.7.
- *3dSkullStrip* is a modified version of BET that is incorporated in the AFNI toolkit [21]. The algorithm begins by preprocessing the image to correct for spatial variations in image intensity and repositioning the brain to roughly the center of the image. Then a modified algorithm based on BET is used to expand a mesh sphere until it envelops the entire brain surface. Among the modifications are procedures to avoid the eyes and ventricles and operations to avoid cutting into the brain. The version of the AFNI toolkit that was used was AFNI_2011_12_21_1014.
- *FreeSurfer’s Hybrid Watershed Algorithm (HWA)* is a hybrid technique that uses a watershed algorithm in combination with a deformable surface algorithm [22]. The watershed algorithm is first used to create an initial mask under the assumption of the connectivity of white matter. Then a deformable surface model is used to incorporate geometric constraints into the mask. The version of FreeSurfer that was used was 5.3.0.

#### Data Analysis

To illustrate the use of the NFBS as testing data, it was used to compare the performance of BET, 3dSkull-Strip and HWA for automatically skull-stripping the original NFB data. In a second analysis we compared the performance of the NFBS BEaST library to the default BEaST library and the three aforementioned methods. Each of the methods were used to skull-strip data from the IBSR (version 2.0) and LPBA40 [3, 4]. To insure consistent image orientation across methods and datasets, they were all converted to LPI orientation using AFNI’s 3dresample program [21]. Additionally, a step function was applied to all of the outputs using AFNI’s 3dcalc tool to binarize all of the generated masks.

The performance of the various methods were compared using the Dice similarity [23] between the mask generated for an image and its corresponding reference (‘gold standard’) mask. Dice was calculated using: *D* = 2 · |*A* ⋂ *B*|/(|*A*| + |*B*|), where *A* is the set of voxels in the test mask, B is the set of voxels in the gold standard data mask, *A* ⋂ *B* is the intersection of *A* and *B*, and | · | is the number of voxels in a set. Dice was implemented in custom Python scripts that used the NiBabel neuroimaging package [24] for data input. Dice coefficients were subsequently graphed as box plots using the ggplot2 package [?] for the R statistical computing language [25].

#### Results

Figure 2 displays box plots of the Dice coefficients that result from using NFBS as gold standard data. The results indicate that 3dSkullStrip performed significantly better of the three methods, with HWA coming in second. In particular, average Dice similarity coefficients were 0.893 ± 0.027 for BET, 0.949 ± 0.009 for 3dSkullStrip, and 0.900 ± 0.011 for HWA. It is perhaps worth noting that BET, the method that performed worst on the NFBS library, took substantially more time to run (25 min) compared to 3dSkullStrip (2 min) and HWA (1 min.).

**Figure 2.**
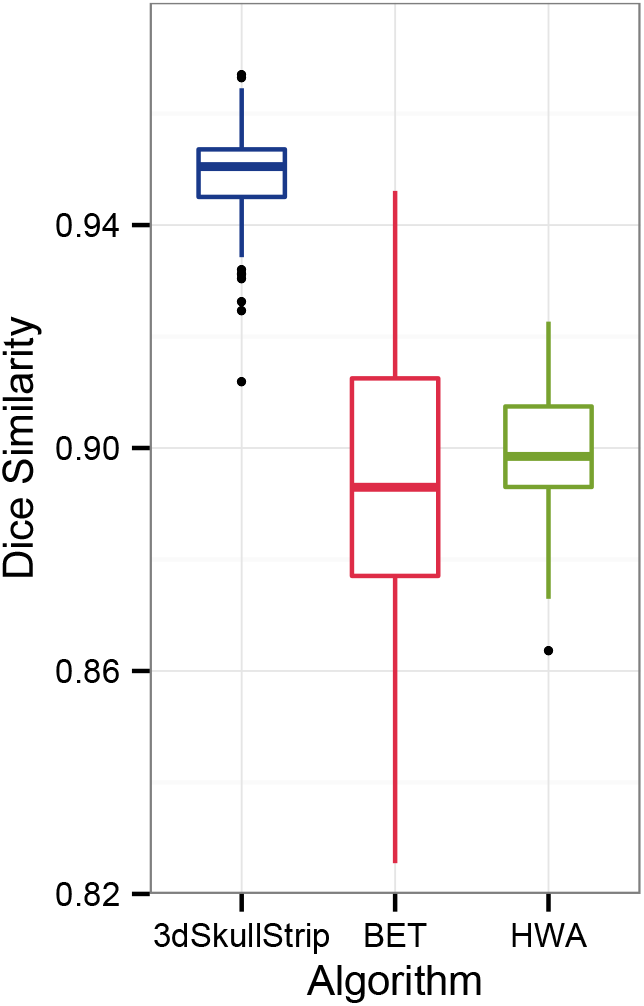
Comparison of methods on NFBS.

Boxplots of Dice coefficients measuring the similarity between masks generated from each image using BET, 3dSkullStrip HWA and the image’s corresponding reference brain masks.

Switching now from using NFBS as the repository of gold standard skull-stripped images to using the IBSR and LPBA40 repositories as the source of gold standard images, Figure 3 shows box plots of the Dice similarity coefficients for BET, 3dSkullStrip, HWA, BEaST using beast-library-1.1, and BEaST using NFBS as the library of priors. For IBSR, 3dSkull-Strip performs better than BET and HWA, similarly to NFBS. However, for LPBA40, BET performs much better than the other two algorithms. The BEaST method was also applied to the anatomical data in these repositories using two different methods: first with the original beast-library-1.1 set as the prior library, and second with the entire NFBS set as the prior library.

**Figure 3.**
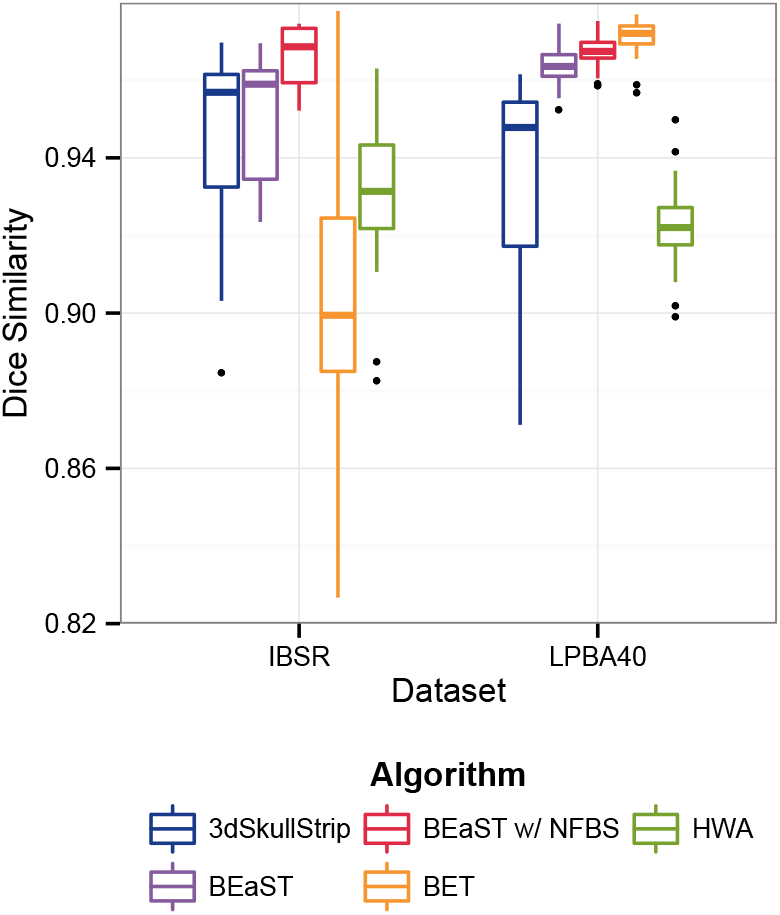
Dice Similarity Coefficients for IBSR & LPBA40.

Box plot of Dice coefficients for BET, 3dSkullStrip, HWA, BEaST using beast-library-1.1, and BEaST using NFBS as the library of priors. One subject was left out of the Dice calculation for each of the following: BEaST w/ beast-library-1.1 on IBSR (IBSR_11), BEaST w/ beast-library-1.1 on LPBA40 (S35), and BEaST w/ NFBS on LPBA40 (S35).

For the BEaST method, it can be said that using NFBS as the prior library resulted in higher average Dice similarity coefficients and smaller standard deviations^[1]^. Differences in Dice coefficients between datasets may be due the size and quality of the NFB study, as well as the pathology and age of the participants. There also may be differences in the standard of the masks, such as length of brainstem and inclusion of exterior nerves and sinuses.

Placing our results in the context of other skull-stripping comparisons, differences between the Dice coefficients reported here and values already published in the literature may be due to the version and implementation of the skull stripping algorithms, a possibility that has received support in the literature [6]. These differences may also result from our application of AFNI’s 3dcalc step function to the skull-stripped images in order to get a value determined more by brain tissue and less influences by CSF. As the NFBS dataset is freely accessible by members of the neuroimaging community, these possibilities may be investigated by the interested researcher.

## Discussion

In summary, we have created and shared the NFBS repository of high quality, skull-stripped T1-weighted anatomical images that is notable for its quality, its heterogeneity, and its ease of access. The procedure used to populate the repository combined the automated, state-of-the-art BEaST algorithm with meticulous hand editing to correct any residual brain extraction errors noticed on visual inspection. The manually corrected brain masks will be a valuable resource for improving the quality of preprocessing obtainable on the NFB data. The corresponding BEaST library will improve skull-stripping of future NFB releases and may outperform the default beast-library-1.1 on other datasets [see Figure 3]. Additionally, the corrected brain masks may be used as gold standards for comparing alternative brain extraction algorithms, as was illustrated in our preliminary analysis [see Figure 2].

The NFBS repository is larger and more heterogeneous than many comparable datasets. It contains 125 skull-stripped images, is composed of images from individuals with ages ranging from 21-45, and represents individuals diagnosed with a wide range of psychiatric disorders [see Table 1]. This variation is a crucial feature of NFBS, as it accounts for more than the average brain. Ultimately, this variation may prove useful for researchers interested in developing and evaluating predictive machine learning algorithms on both normal populations and those with brain disorders [26].

Finally, the repository is completely open to the neuroscience community. NFBS contains no sensitive personal health information, so researchers interested in using it may do so without submitting an application or signing a data usage agreement. This is in contrast to datasets such as the one collected by the Alzheimer’s Disease Neuroimaging Initiative (ADNI) [1]. Researchers can use ADNI to develop and test skull-stripping algorithms [18], but in order to do so must first apply and sign a data usage agreement, which bars them from distributing the results of their efforts. Thus, we feel that NFBS has the potential to accelerate the pace of discovery in the field, a view that resonates with perspectives on the importance of making neuroimaging repositories easy to access and easy to use [27].

## Availability of supporting data

The NFBS skull-stripped repository is available at: https://preprocessed-connectomes-project.org/NFB_skullstripped. Bash and Python scripts used for this paper are available on GitHub at: https://github.com/preprocessed-connectomes-project/NFB_skullstripped.

## Abbreviations

MRI: : magnetic resonance imaging;
NFBS: : Neurofeedback Skull-stripped;
CSF: : cerebrospinal fluid;
BEaST: : brain extraction based on nonlocal segmentation technique;
BET: : brain extraction technique;
HWA: : hybrid watershed technique;
IBSR: : Internet brain segmentation repository;
LPBA40: : LONI Probabilistic Brain Atlas;
ADNI: : Alzheimer’s Disease Neuroimaging Initiative

## Author’s contributions

RCC designed the Neurofeedback study and Skull-stripped repository; BP and EST performed manual correction and validation of results; BP performed the validation analyses; BP, RCC, JSP, and JPP wrote the data note. All authors read and approved of the final version.

## Acknowledgements

We would like to thank Dr. Simon Fristed Eskildsen for help with the installation and optimization of the BEaST method. We would also like to acknowledge Qingyang Li for creating the BEaST guide, as well as the Bash script that we based our script on. Lastly, we would like to thank all of those involved in the participation, data collection, and data sharing initiative of the Enhanced Rockland Sample. This work was supported by R01MH101555 from the National Institute of Mental Health to RCC.

## Footnotes

[1] BEaST was unable to segment 1 subject, IBSR_11, in IBSR, only when using beast-library-1.1. For LPBA40, BEaST was also unable to segment 1 subject, S35, when using beast-library-1.1 and NFBS. These subjects were left out of the Dice calculations.

